# *Ralstonia solanacearum* pandemic lineage strain UW551 overcomes inhibitory xylem chemistry to break tomato bacterial wilt resistance

**DOI:** 10.1101/2023.01.20.523839

**Authors:** Corri D. Hamilton, Beatriz Zaricor, Carolyn Jean Dye, Emma Dresserl, Renee Michaels, Caitilyn Allen

**Author notes:** Department of Microbiology and Immunology, University of British Columbia, Vancouver, Canada (Corri D. Hamilton). **Author contributions:** CDH designed experiments, conducted experiments, analyzed data, and wrote the manuscript; BZ, ED, and CD conducted experiments; CA designed experiments and wrote the manuscript.

## Abstract

Plant pathogenic *Ralstonia* strains cause bacterial wilt disease by colonizing xylem vessels of many crops, including tomato. Host resistance is the best control for bacterial wilt, but resistance mechanisms of the widely used Hawaii7996 tomato breeding line are unknown. Using growth in *ex vivo* xylem sap as a proxy for host xylem, we found that *Ralstonia* strain GMI1000 grows in sap from both healthy plants and *Ralstonia*-infected susceptible plants. However, sap from *Ralstonia*-infected Hawaii7996 plants inhibited *Ralstonia* growth, suggesting that in response to *Ralstonia* infection, resistant plants increase inhibitors in their xylem sap. Consistent with this, reciprocal grafting and defense gene expression experiments indicated that Hawaii7996 wilt resistance acts both above- and belowground. Concerningly, Hawaii7996 resistance is broken by *Ralstonia* strain UW551 of the pandemic lineage that threatens highland tropical agriculture. Unlike other *Ralstonia* strains, UW551 grew well in sap from *Ralstonia*-infected Hawaii7996 plants. Moreover, other *Ralstonia* strains could grow in sap from Hawaii7996 plants previously infected by UW551. Thus, UW551 overcomes Hawaii7996 resistance in part by detoxifying inhibitors in xylem sap. Testing a panel of xylem sap compounds identified by metabolomics revealed that no single chemical differentially inhibits *Ralstonia* strains that cannot infect Hawaii7996. However, sap from *Ralstonia*-infected Hawaii7996 contained more phenolic compounds, which are known plant antimicrobial defenses. Culturing UW551 in this sap reduced total phenolic levels, indicating that the resistance-breaking *Ralstonia* strain degrades these chemical defenses. Together, these results suggest that Hawaii7996 tomato wilt resistance depends at least in part on inducible phenolic compounds in xylem sap.

## 1. Introduction

Tomato (*Solanum lycopersicum*), an important source of nutrients like vitamins A and C, antioxidants, and carotenoids, is widely cultivated, with global production exceeding 70 million metric tons each year (García-Alonso et al. 2009; Perveen et al. 2015). Tomatoes are the leading processing vegetable crop in the United States with a value of $1.6 billion in 2020 (NASS USDA 2020). However, a plethora of diseases reduce tomato production. Soilborne bacteria in the *Ralstonia solanacearum* species complex (RSSC), which cause bacterial wilt disease, are among the most damaging tomato pathogens (Hayward et al. 1994; Mansfield et al. 2012). Controlling bacterial wilt losses is important for tomato farmers at every scale from subsistence plots to large-scale industrial plantations (Elphinstone. 2005).

The most effective and environmentally friendly way to manage bacterial wilt is to plant crop varieties that are genetically resistant to the pathogen (Boshou 2008; Lebeau et al. 2011). Disease losses are not reliably reduced by cultural or even harsh chemical measures (Enfinger et al. 1979; Gan et al. 2000). The lack of other effective management strategies increases the importance of breeding tomatoes for bacterial wilt resistance (Scott et al. 2005; Hong et al. 2008; Yuliar & Toyota, 2015). There are three major resistance sources (Hawaii, CRA 66, and PI129080), all derived from wild relatives of small-fruited tomato species (Gilbert et al. 1973; Prior et al. 1994; Henderson & Jenkins, 1972; Grimault and Prior, 1993; Grimault et al., 1995). The standouts among these sources are the highly resistant ‘Hawaii’ breeding lines: H7996, H7997 and H7998, which were probably derived from crosses between *S. lycopersicum* and *S. pimpinellifolium* (Gilbert et al., 1973). An evaluation of over 35 tomato wilt resistance sources in 11 countries found that H7996 confers the most stable and effective bacterial wilt resistance (Wang et al 1998). Hawaii lines have several quantitative trait loci (QTL) that mediate wilt resistance. Introgressing this resistance into commercial cultivars is difficult because undesirable agronomic traits like small fruit size are tightly linked to the resistance QTLs (Opina et al., 1990; Wang et al. 1998; Carmeille et al. 2006; Yuliar & Toyota, 2015). Despite these difficulties, the Hawaii breeding lines are the source of nearly all tomato bacterial wilt resistance worldwide (Lopes et al. 2022; Brand-Daunay et al. 2010; Ho et al. 2013; Scott et al. 2009; Wang et al. 2000).

Plant pathogenic *Ralstonia* cause bacterial wilt by entering host roots from the soil, moving to the developing vascular bundles and spreading systemically through the plant’s water-transporting xylem vessels (Caldwell et al. 2017; McGarvey et al. 1999). The pathogen rapidly grows to high populations *in planta*, producing virulence factors including plant cell wall-degrading enzymes, many type III-secreted effectors, and extracellular polysaccharide (EPS), leading to water transport obstruction, characteristic wilting symptoms, and plant death (Denny 2006; Lowe-Power et al., 2018b). *Ralstonia* cycles between two very different physiochemical environments: rhizosphere soil and host xylem sap. The 12 known bacterial wilt resistance QTL have been mapped, but we lack functional understanding of wilt resistance (Thoquet et al., 1996; Scott et al., 2005; Wang et al., 2013). Studies have focused primarily on belowground resistance mechanisms, such as the role of root architecture in bacterial entry (Nakaho et al., 2004; Caldwell et al., 2017). However, H7996 resistance mechanisms include upregulation of plant defense signaling pathways in stems and restricting (but not eliminating) stem colonization (Milling 2011). Grafting experiments confirmed that even though *Ralstonia* aggressively colonizes the roots of susceptible varieties, grafting H7996 scions onto susceptible rootstocks limits pathogen movement up into plant stems (Planas-Marques et al 2020). Additionally, plant defensive reactive oxygen species increase differentially in resistant stems during infection with wild-type *Ralstonia* compared to a mutant missing the major virulence factor EPS (Milling et al., 2011; Jones and Dangl, 2006). Further, *Ralstonia* alters the xylem environment during infection of susceptible tomato cultivars (Lowe-Power et al., 2018; Lowe-Power et al., 2018b; Jacobs et. al., 2012). Together, these findings led us to ask if the xylem environment plays a role in bacterial wilt resistance.

The wilt resistance in line H7996 has been generally durable, but it is not infallible. A few strains of *Ralstonia* can break this resistance to cause bacterial wilt on H7996 (Wang et al 1998; Jaunet and Wang 1999). We found that one such strain is UW551, representative of a near-clonal pandemic lineage that originated in the South American Andes and causes both tomato bacterial wilt and potato brown rot disease in cool tropical highlands and temperate zones (Clarke et al., 2015 Phytopathology). Agricultural trade has spread this pandemic lineage around the world, displacing endemic strains and causing major losses to potato, a key carbohydrate crop (Ravelomanantsoa et al., 2018; Sharma et al., 2021; Sharma et al., 2022). The pandemic lineage, known historically and for regulatory purposes as Race 3 biovar 2, is a U.S. Select Agent pathogen (Lambert, 2002; Hayes et al., 2022). Understanding the mechanism of host resistance will be key to developing effective disease resistance to mitigate this threat (Kim et al. 2016).

The concerning discovery that the *Ralstonia* pandemic lineage can wilt H7996 tomatoes increases the urgency of understanding bacterial wilt resistance mechanisms. We found that in response to infection by *Ralstonia* strain GMI1000, the xylem sap of wilt resistant H7996 tomato plants becomes toxic and restricts GMI1000 growth. In contrast, resistance-breaking strain UW551 grows well in this sap. We separately tested the hypotheses that H7996 chemical defenses were overcome by: 1) subverting host defense; 2) increasing bacterial toxin efflux: or 3) chemical degradation. We used metabolomics, targeted mutants, and bacterial growth assays to determine that *Ralstonia* UW551 succeeds in xylem of resistant hosts by degrading host defensive phenolic compounds.

## 2. Results

### 2.1 Ralstonia strain UW551 overcomes Hawaii7996 wilt resistance

The H7996 breeding line is the primary source of resistance to tomato bacterial wilt, but this resistance has occasionally failed in tropical highlands where the *Ralstonia* potato brown rot pandemic lineage is endemic (Philippe Prior, personal communication). To experimentally determine if the pandemic lineage can break H7996 resistance, we directly measured the virulence of the typical pandemic lineage strain UW551 on H7996 using a naturalistic soil soak inoculation under controlled growth chamber conditions. As expected, phylotype I strain GMI1000 did not wilt H7996 plants, although both GMI1000 and UW551 were fully virulent on susceptible tomato cv. Money Maker (MM) (Fig 1a). GMI1000 was rarely (<8%) detected in the xylem of inoculated H7996 plants, but this strain never caused wilt symptoms on H7996 (data not shown; Fig 1a). In contrast, pandemic lineage strain UW551 killed 42% of inoculated H7996 plants and the pathogen was detected in more than half the asymptomatic plants (Fig 1b). These results demonstrated that H7996 has little resistance to *Ralstonia* strain UW551.

**Figure 1.**
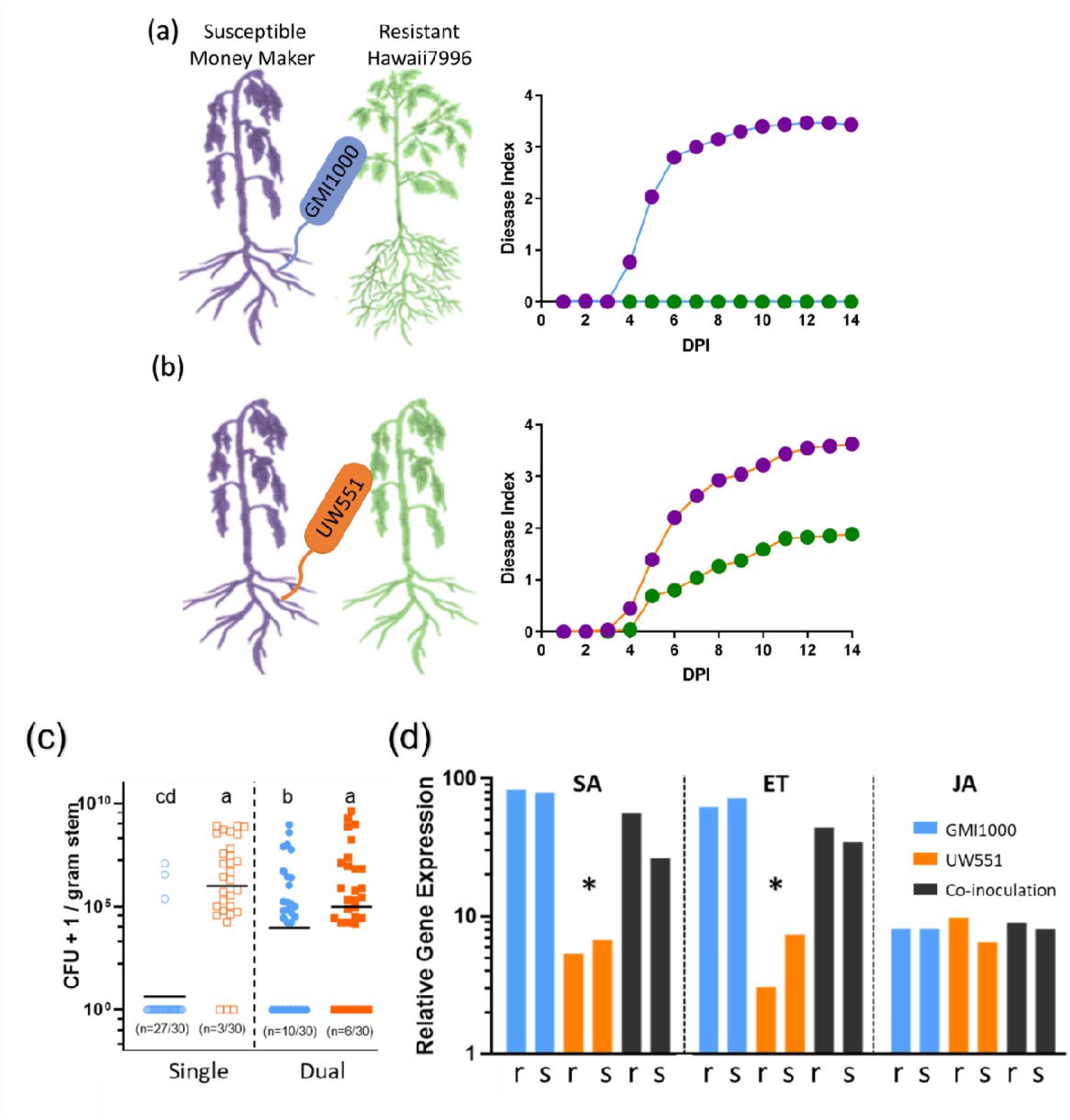
*Ralstonia* strains UW551 and GMI1000 differ in virulence and induction of defense responses on wilt-susceptible and wilt-resistant tomato plants. (**a, b**) Disease progress of *Ralstonia* GMI1000 (blue line) or *Ralstonia* UW551 (orange line) on susceptible host Money Maker (purple circles) or resistant host Hawaii7996 (green circles) was rated on a 0 to 4 disease index scale. (**c**) Unwounded 17-day-old tomato plants were inoculated with *Ralstonia* GMI1000 (open blue), *Ralstonia* UW551 (open orange), or a 1:1 mixture of both strains (closed symbols) and bacterial population size was determined after 5 days. Horizontal bars indicate geometric mean; letters indicate differences at P<0.05, Mann-Whitney Test. (**d**) Relative expression levels of tomato SA, ET, and JA defense pathway were measured by qRT-PCR in roots (r) or stems (s) of whole H7996 tomato plants 96 h after inoculation with either *Ralstonia* strain GMI1000 (blue bar), UW551 (orange bars), or co-inoculation with both strains (black bars). Gene expression levels are relative to those in water-inoculated healthy control plants. Asterisks indicate difference at P<0.05, T-test. All plants were drench-inoculated to ∼ 5×10^7^ CFU/g of soil. Experiments contain 3 biological assays with a total of 9-15 plants.

### 2.2 Pandemic lineage strain *Ralstonia* UW551 evades Hawaii7996 plant defenses

To isolate the effects of wilt-resistant xylem on *Ralstonia* from resistance acting in other tissues, we measured symptoms and bacterial colonization of H7996 plants following petiole inoculation, which bypasses the roots entirely. *Ralstonia* GMI1000 colonized the stems of resistant H7996 stems only 10% of the time and never caused symptoms. In contrast, strain UW551 colonized 90% of H7996 stems and many developed symptoms (Fig 1c). Co-inoculating plants with a 1:1 mixture of GMI1000 and UW551 helped GMI1000 colonize H7996, with 33% of co-inoculated stems containing GMI1000 while UW551 was detected in 80% of co-inoculated stems. This is further evidence that conditions in stems of H7996 contribute to this tomato breeding line’s wilt resistance (Fig 1c).

To understand why co-inoculation slightly reduced UW551 colonization, we compared the transcriptional responses of resistant plants to single and co-inoculation with *Ralstonia* strains. Tomato PR1-a and PR1-b genes are reporters of the salicylic acid (SA) and both the SA and the ethylene (ET) defense pathways, respectively. Both genes are up-regulated in response to *Ralstonia* infection (Milling 2011). The jasmonic acid (JA) pathway reporter gene PIN2, which is not induced by *Ralstonia* under these conditions, was included as a negative control. Strain GMI1000 triggered strongly increased expression of both PR1-a and PR1-b. Regardless of host tissue (roots or stems), UW551 triggered a smaller upregulation of the SA and ET pathways relative to plants infected with GMI1000 alone or with GMI1000 and UW551 together (Fig 1d). The fact that resistance-breaking strain UW551 is confronted with a full defense response when it co-infects with GMI1000 suggests that UW551 somehow evades recognition by H7996 plants rather than suppressing defenses.

### 2.3 Bacterial wilt resistance in H7996 acts in aboveground plant parts as well as in roots

Grafting wilt-susceptible tomato scions onto H7996 rootstock can control bacterial wilt because H7996 restricts movement of *Ralstonia* from the roots up into the stem (Cardoso et al. 2012; Planas-Marques et al 2020). However, although bacterial virulence correlates with stem colonization, many colonized plants are asymptomatic. Under our conditions, chimeric plants consisting of H7996 rootstock grafted to wilt-susceptible MM tomato were fully resistant to *Ralstonia* strain GMI1000 (Fig 2a). The reciprocal composite plants, which consisted of susceptible rootstock grafted onto the resistant scion, were sometimes wilted by GMI1000 but were much less susceptible than MM plants (Fig 2a). This indicates that an element of H7996 wilt resistance acts in aboveground parts of the plant, where *Ralstonia* is largely confined to the water-transporting xylem vessels.

**Figure 2:**
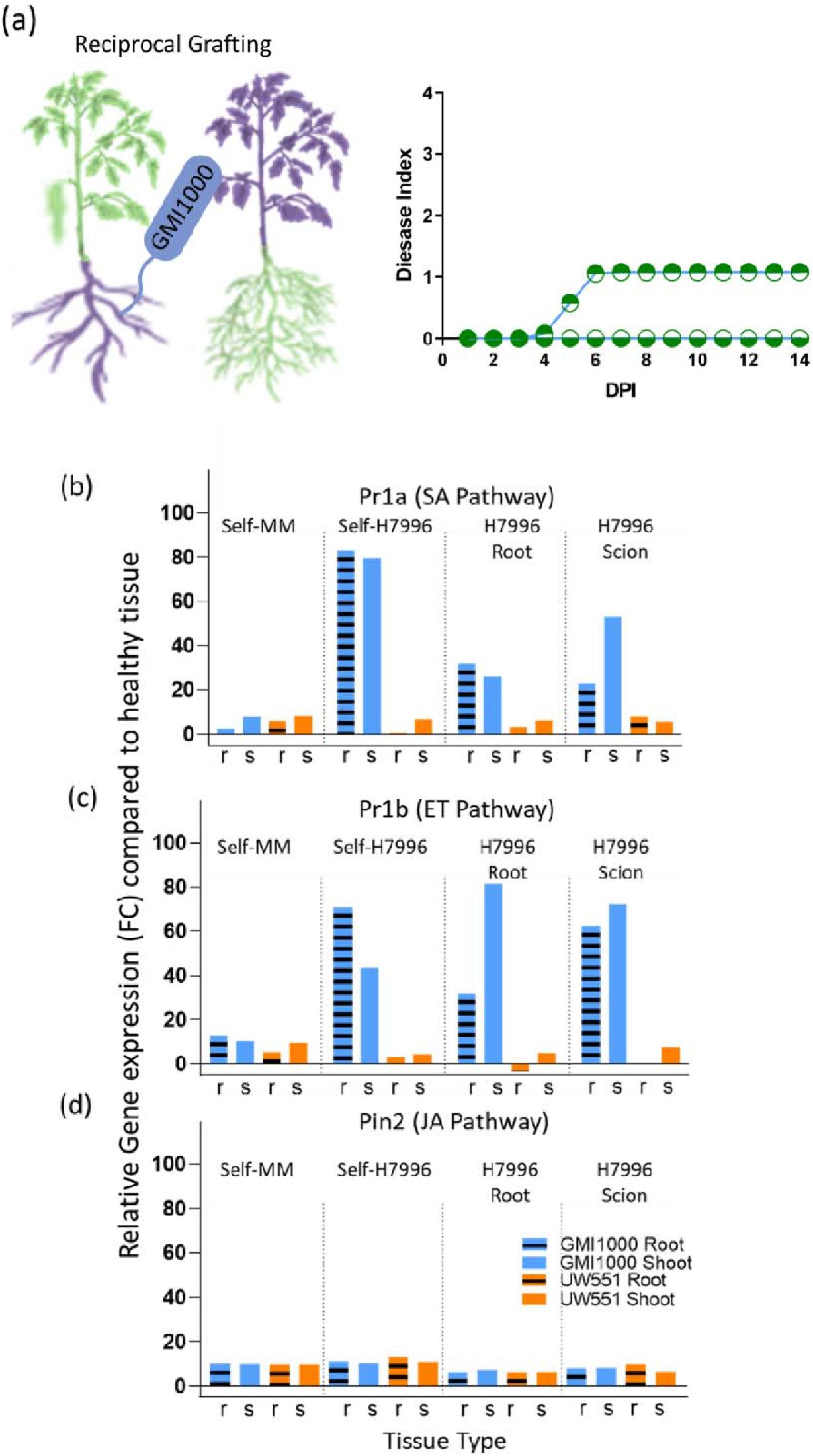
Grafting bacterial wilt*-*resistant H7996 scions to susceptible rootstocks increases plant defense gene expression in roots. (**a**) Disease progress of *Ralstonia* GMI1000 (blue line) on reciprocally grafted Money Maker (green)-H7996 (purple) chimeric plants. Green circles with open bottoms indicate mean disease index of susceptible Money Maker rootstocks grafted to resistant Hawaii7996 scions while green circles with open tops indicate mean disease index of resistant H7996 rootstock grafted to susceptible Money Maker scions. (**b-d**) Relative expression levels of tomato SA, ET, and JA defense pathway reporter genes were measured by qRT-PCR in H7996 roots (r; dotted bar) or stems (s; solid bar) of grafted tomato plants 96 h after soil-soak inoculation with either *Ralstonia* GMI1000 (blue bars) or *Ralstonia* UW551 (orange bars). Gene expression levels are relative to those in healthy water-inoculated control plants. Data are shown as mean fold-change expression of 3 independent experiments; asterisks indicate significant differences from uninoculated control (P<.05, T-test).

To better understand the link between xylem tissue and wilt disease suppression, we measured expression of tomato defense genes in roots or stems following inoculation of grafted plants with either *Ralstonia* strain GMI1000, which cannot wilt H7996, or UW551, which overcomes H7996 wilt resistance. Tomato plants of each genotype were grafted to themselves to control for any effects of the grafting procedure. Relative to healthy control plants, *Ralstonia* infection triggered increased expression of SA defense pathway marker PR1-a regardless of host susceptibility or tissue. Wilt-susceptible MM responded similarly to both *Ralstonia* strains, but in wilt-resistant H7996 plants, strain GMI1000 triggered higher PR1-a expression than UW551; this was true in both stems and roots (Fig. 2b). In reciprocally grafted tomato plants, PR1-a transcription increased in roots and stems not only when the rootstock was H7996, as expected, but also when the rootstock was susceptible MM and the only resistant tissue was in the H7996-derived stems. Similar trends were seen for expression of SA/ET marker PR1b, and, to a lesser degree, of JA pathway marker PIN2 (Fig. 2c-d). Taken together, the reciprocal grafting experiments show that H7996 wilt resistance does not only act belowground. Grafting a resistant scion onto susceptible rootstock decreased soilborne wilt disease incidence and increased defense expression even in susceptible root tissue. Thus, to succeed in a resistant plant, *Ralstonia* must overcome defenses in both roots and stems.

### 2.4 H7996 xylem sap inhibits growth of *Ralstonia* strains that cannot break resistance

We previously observed that infection by *Ralstonia* strain GMI1000 made *ex vivo* xylem sap from wilt-susceptible Bonny Best tomato plants into a better bacterial growth medium than sap from healthy plants. Metabolomic analysis showed that *Ralstonia* infection increases levels of several carbon and nitrogen sources in xylem sap (Lowe-Power et al. 2018). We obtained a similar result using sap from Money Maker (MM), another wilt-susceptible cultivar (Supplemental Fig S1a). We wondered if the ability to break H7996 resistance correlates with the ability of an *Ralstonia* strain to grow in H7996 xylem sap. Strains UW551 and GMI1000 grew equally well on sap from healthy H7996 tomato plants (Fig 3a). However, strain GMI1000, which cannot wilt H7996, grew very poorly on sap from H7996 plants that had been infected with GMI1000 (*P*<.001, T-test) (Fig 3a). In striking contrast, resistance-breaking strain UW551 actually grew better on *ex vivo* xylem sap from H7996 plants previously infected by UW551 (*P*<.05, T-test) (Fig 3a). Interestingly, xylem sap from H7996 plants that were previously infected by UW551 was a good growth medium for strain GMI1000.

**Figure 3.**
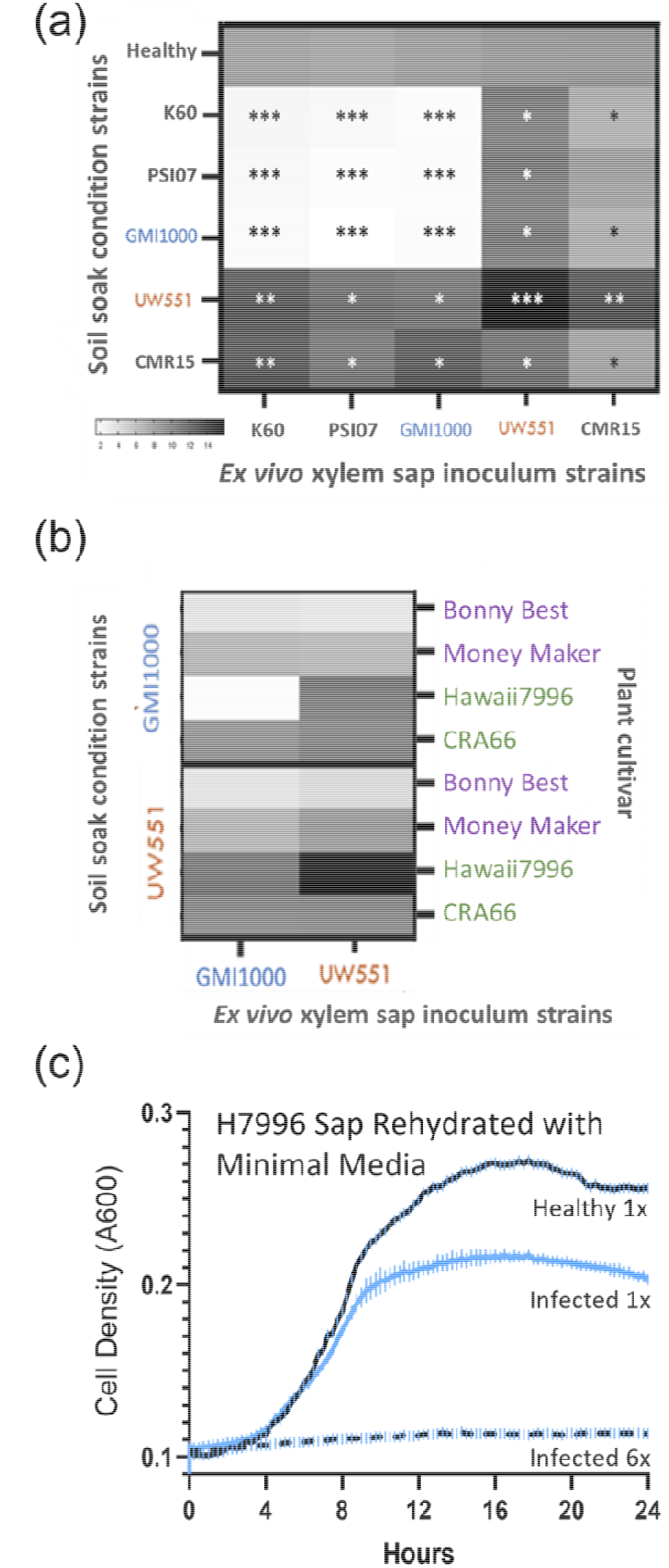
Resistance-breaking UW551 overcomes the inhibitory xylem conditions formed during infection of H7996. (**a**) Growth of diverse *Ralstonia* strains (indicated on X axis) on *ex vivo* xylem sap from H7996 plants that were healthy or soil-soak inoculated by diverse *Ralstonia* strains (indicated on the Y-axis). Growth was quantified as area under a 24-hour growth curve indicated by shading from white (low area or little to no growth) to black (maximum growth). Asterisks indicate significant difference from bacterial growth in sap from healthy plants, determined by T-test, *** P<.001, **P <. 001, ** P<.05. (**b**) Growth of *Ralstonia* GMI1000 and UW551 (indicated on X axis) on *ex vivo* xylem sap harvested from Bonny Best (susceptible), Money Maker (susceptible), and CRA66 (resistant) tomatoes that were infected by GMI1000 or UW551 (indicated on the Y-axis). Growth was quantified as area under the growth curve as for (b) above. For all *ex vivo* sap experiments, bacterial growth was measured over 24 h spectrophotometrically as A_600nm_ in a plate reader. (**c**) Growth of *Ralstonia* strain GMI1000 in *ex vivo* xylem sap harvested from H7996 tomato plants that were healthy (black line) or infected by GMI1000 (blue lines). Xylem sap was filter-sterilized, dehydrated and rehydrated with minimal medium to 1X original concentration (solid lines), or 6X original concentration (dashed line). Data are shown as means of 3 independent experiments, each containing 3 technical replicates.

The RSSC comprises three species and four geographical origin lineages called phylotypes (Safni et al., 2014; Prior and Fegan, 2005). To see if our results were generalizable across the RSSC, we measured xylem sap growth of tomato-infecting strains from all four phylotypes. These were strain PSI07 from phylotype IV (*R. syzygii*), strains GMI1000 and CMR15 from phylotype I (*R. pseudosolanacearum)*, and strains K60 and UW551 from phylotype IIA and IIB, respectively (*R. solanacearum)* (Poussier et al., 2000; Fegan and Prior, 2005; Jeong et al., 2007; Sharma et al., 2022). These five diverse *Ralstonia* strains grew equally well on sap from healthy H7996 tomato plants (Fig 3b). However, *ex vivo* sap harvested from plants infected by a strain that cannot break H7996 resistance inhibited growth of all *Ralstonia* strains that cannot break H7996 resistance. Like GMI1000, strains PSI07 and K60 cannot break H7996 resistance. GMI1000, PSI07 and K60 barely grew in *ex vivo* xylem sap from H7996 plants infected by any of the three strains (P<.001 relative to growth in sap from healthy plants, T-test) (Fig 3a; Supplemental Fig S1b). In contrast, *Ralstonia* strains UW551 and CMR15, which can both break H7996 resistance, grew better in *ex vivo* xylem sap from H7996 plants that had been infected by either resistance-breaking strain than in sap from healthy plants (*P*<.05 T-test) (Fig 3b; Supplemental Fig S1a). Together, these *ex vivo* growth experiments suggested that UW551 infection somehow conditions the xylem sap of H7996 plants to make it a good growth medium for any *Ralstonia* strain regardless of whether it can break resistance.

### 2.5 Inhibitory xylem sap is specific to the wilt resistance in H7996

To determine if xylem sap growth inhibition is a unique property of H7996, we measured growth of the five *Ralstonia* strains on *ex vivo* xylem sap harvested from diverse tomato genotypes. These included wilt-susceptible tomato varieties Bonny Best and Money Maker and wilt-resistant tomato lines H7996 and CRA66. The various *Ralstonia* strains grew similarly on sap from the two susceptible tomato cultivars (Fig 3b; Supplemental Fig S1a). However, xylem sap from wilt-resistant CRA66 plants that had been conditioned by previous *Ralstonia* infection supported bacterial growth as well as sap from healthy CRA66 plants (*P*>.05) (Fig 3b; Sup. Fig S1a). This suggests that unlike H7996, wilt resistance of CRA66 tomato does not involve inhibiting bacterial growth in xylem sap.

### 2.6 Poor growth of GMI1000 in H7996 xylem sap is not explained by nutrient limitation

Pathogenic bacteria sometimes cause plants to relocate nutrients to support bacterial growth; conversely, resistant plants can sequester nutrients to slow pathogen growth. To test the hypothesis that strain GMI1000 grows poorly in sap from infected H7996 plants because it contains fewer nutrients, we dehydrated pooled xylem sap from GMI1000-infected H7996 plants and rehydrated it in minimal medium with no carbon source at either its original volume or 1/6 the original volume. As expected, strain GMI1000 grew less in the full-strength rehydrated sap from the infected H7996 plants than in sap from healthy plants. But rather than increasing nutrients, concentrating this sap six-fold completely inhibited bacterial growth (RM-ANOVA, P<.05) (Fig 3c). We concluded that the poor growth of GMI1000 in H7996 sap was not caused by starvation but rather by the presence of inhibitors.

### 2.7 Resistance-breaking *Ralstonia* UW551 facilitates subsequent infection of Hawaii7996 stems by GMI1000

To better understand how resistance-breaking *Ralstonia* strain UW551 overcomes inhibitors in H7996 xylem sap, we asked if exposing *ex vivo* xylem sap to UW551 could make it less toxic to strain GMI1000. We harvested xylem sap from GMI1000-infected H7996 plants, pooled and filter-sterilized it, then used it as a growth medium for either GMI1000 or UW551. This “conditioned” sap was again pooled and filter-sterilized, supplemented with 20 mM sucrose to ensure adequate carbon, and then used it as a growth medium for strain GMI1000. Although GMI1000 did not thrive in either of these conditioned media, it grew significantly better in the sap that previously hosted resistance-breaking strain UW551. This suggested that UW551 inactivated antibacterial compounds that H7996 produced in response to GMI1000 infection (Fig 4a).

**Figure 4.**
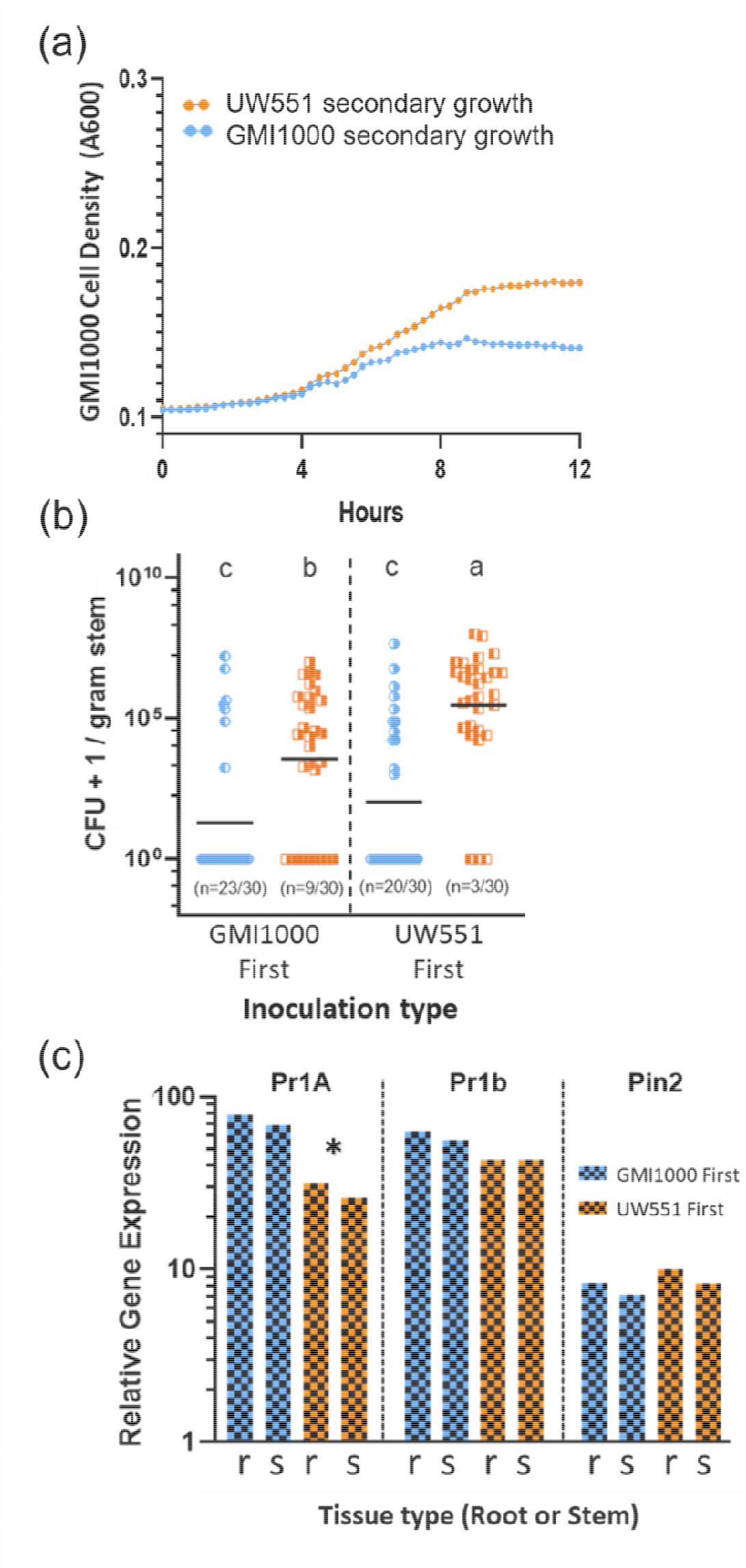
Resistance-breaking *Ralstonia* UW551 alters the Hawaii7996 stem environment to favor GMI1000. (**a**) Growth of *Ralstonia* GMI1000 in *ex vivo* xylem sap that was harvested from GMI1000-infected H7996 plants, used as a growth medium for GMI1000 (blue dots) or UW551 (orange dots), and then filter-sterilized and supplemented with 20 mM sucrose to ensure adequate carbon source. (**b**) *Ralstonia* GMI1000 and UW551 colonization of H7996 tomato stems sampled 4 days after plants were soil-soak inoculated with GMI1000 first, then with UW551 24 h later (left box) or in the reverse order (right box). Each dot indicates the bacterial population size in one plant; N=30 plants for each treatment; the number of plants with undetectable populations is given in parentheses below each column; horizontal bar indicates geometric mean population. Letters above each column indicate differences at P< .05, t-test. (**c**) Relative expression levels of SA, ET, and JA defense pathway marker genes were measured by qRT-PCR in tomato roots (r) or stems (s) at 96 h after soil-soak inoculation with either *Ralstonia* strain GMI1000 first (blue) or UW551 first (orange) as described above. Gene expression levels are relative to those in water-inoculated healthy control plants. Data are shown as mean expression levels from 3 independent experiments, each containing RNA from 12 plant samples; asterisk indicates significant difference between treatments (P<.05, T-test).

We tested this hypothesis in a more biologically relevant context by soil inoculating H7996 plants with either GMI1000, followed 24 h later by a second inoculation with UW551, or in the opposite order. In both conditions, after 4 days UW551 reached a higher mean population size in stems than GMI1000 and was more likely than GMI1000 to reach detectable levels in stems. However, when resistant plants were exposed first to UW551, GMI1000 was detectable in 34% of plants compared to only 24% when GMI1000 was inoculated first. The mean GMI1000 stem population size trended higher when UW551 was inoculated first but was not significantly different (Fig 4b). When H7996 plants were exposed to GMI1000 first, mean stem population size of UW551 was lower than in plants initially inoculated with UW551 (Fig 4b). This result suggested GMI1000 induced plant defenses that suppressed subsequent colonization by UW551. We tested this hypothesis by comparing expression of H7996 defense marker genes in root and stem tissue from this experiment. In both tissues, initial inoculation with GMI1000 triggered higher expression of SA-pathway marker PR1a (Fig 4c). Expression of Pr1b, associated with both SA and ET defenses, trended lower when UW551 was inoculated first, but the difference was not significant. Taken together, these results show that the mechanisms by which UW551 breaks H7996 resistance involve both suppression of plant defense responses at the transcriptional level and inactivation of antimicrobial chemicals in xylem sap after they are produced.

### 2.8 Xylem sap growth inhibition is not explained by differential tomato glucanase expression

Correlative data suggested that differentially increased β-1,3-glucanases might explain bacterial wilt resistance (Ishihara et al., 2012). To determine if β-1,3-glucanase expression contributes to xylem sap growth inhibition, we compared transcription levels of 3 previously-identified classes of β-1,3-glucanase genes in MM and H7996 stems after inoculation with *Ralstonia* UW551, GMI1000, or water (mock). These experiments confirmed the previous observation that 48 h after inoculation, resistant H7996 tomatoes had much higher β-1,3-glucanase expression than susceptible MM (Fig 4A). This response subsided by 120 h after inoculation (Fig 4B). However, pathogen strain did not affect this transcriptional response; H7996 upregulated its β-1,3-glucanase genes equally in response to GMI1000 and UW551 (Fig 4A). We conclude that differential expression of β-1,3-glucanase genes cannot explain why xylem sap from H7996 inhibits *Ralstonia* GMI1000 but not resistance-breaking strain UW551.

### 2.9 The AcrA and DinF toxin efflux pumps do not explain the ability of UW551 to grow in xylem sap from H7996

Bacteria can mitigate effects of inhibitory compounds by exporting them with toxin efflux pumps. The AcrA and DinF efflux pumps contribute to virulence of *Ralstonia* strain K60 on Bonny Best tomato (Brown at al 2007). We tested the hypothesis that these efflux pumps help *Ralstonia* strain UW551 grow better on H7996 xylem sap by constructing _Δ_*acrA* and _Δ_*dinF* deletion mutants in both GMI1000 and UW551. We used a naturalistic soil soak inoculation assay to measure the virulence of these mutants on susceptible MM tomato, and also measured mutant growth in *ex vivo* xylem sap from infected H7996 plants. As observed in strain K60, _Δ_*acrA* and _Δ_*dinF* mutants in GMI1000 and UW551 were significantly reduced in virulence compared to their respective wildtype parents (Fig 4C and 4D). However, all mutants grew as well as their parent strains on sap from infected and healthy H7996 plants, measured as area under the growth curve. A more sensitive analysis measuring the doubling time of resistance-breaking strain UW551 found no difference between the mutants and the UW551 parent in sap from susceptible tomato regardless of the previously infecting strain (Fig 4E). However, UW551 mutant doubling times in infected H7996 sap were significantly longer than in healthy plant sap whether the plant was previously infected by GMI1000 or UW551 (adjusted P<0.05 by ANOVA with Tukey’s multiple comparisons). This showed that although these toxin pumps impact bacterial growth in H7996 xylem sap, they cannot explain the differential ability of UW551 to grow on *ex vivo* xylem sap from infected H7996.

### 2.10 Physical characterization of xylem sap inhibitors indicated multiple compound types of interest

The observation that culturing UW551 in xylem sap from infected H7996 plants rendered the sap hospitable to subsequent GMI1000 growth strongly suggested that UW551 succeeds in H7996 plants in part because it can detoxify or degrade inhibitory compounds in xylem sap. We subjected *ex vivo* xylem sap to physical and chemical treatment to see if they could abolish the sap’s inhibitory phenotype. The sap still inhibited growth after evaporative dehydration or freezing at -80°C. Inhibitory activity was reduced or eliminated after 15 minutes at 65°C, extraction with phenol:chloroform phase-lock tubes, or treatment with the broad-spectrum Proteinase K. Inhibitory compounds were present in both the eluate and the retentate of a 20 KDa exclusion filter (Supplemental Table S2). Together, these results indicated that the bacterial growth inhibitors in H7996 xylem sap are heat intolerant, proteinaceous, organic, non-volatile aromatics, and include compounds both smaller and larger than 20kDa. These characteristics, some of which are mutually exclusive, correspond to multiple compound classes including known host defense compounds.

### 2.11 Xylem sap metabolite profile of H7996 plants differs in response to infection by GMI1000 vs. UW551

To better understand why H7996 xylem sap inhibits *Ralstonia* growth, we performed a GC-MS metabolomics analysis of *ex vivo* xylem sap from healthy plants and plants previously infected by GMI1000 or resistance-breaking strain UW551. Unsupervised hierarchical clustering and principal component analysis of the results clearly separated the GMI1000 conditioned samples from the UW551 conditioned samples, which were more similar to samples from healthy H7996 plants (Fig 5A, Sup. Fig 2B). Using a cutoff ofLJ≤LJ0.05 (T-test), 81 metabolites were differentially present in xylem sap from *Ralstonia*-infected plants relative to sap from healthy plants (Supplemental Table S1). A subgroup of 22 metabolites were differentially present in sap from plants infected with UW551 compared to those infected with GMI1000 (Fig 5B). Consistent with previous metabolomic analysis of *Ralstonia*-infected Bonny Best sap, sap from infected resistant host H7996 contained higher levels of carbon sources like sucrose relative to sap from healthy plants (*P* < .01) to infected Money Maker (*P* < .05) (Supplemental Table S1).

**Figure 5.**
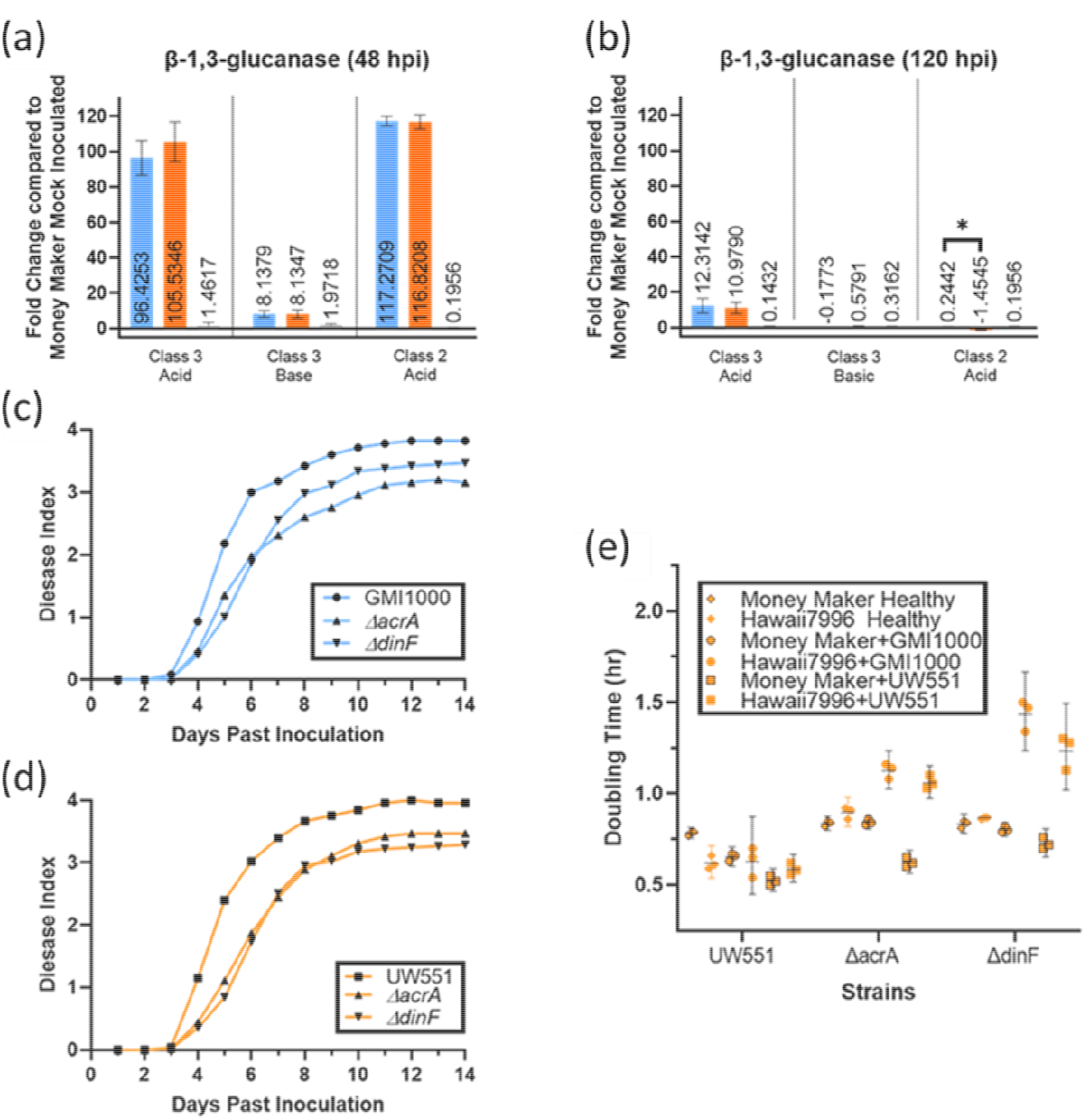
Neither β-1,3 glucanases nor the AcrA or DinF efflux pumps explain why *Ralstonia* strain UW551 breaks resistance or grows better in H7996 xylem sap. Expression of β-1,3-glucanase genes in stems of wilt-resistant H7996 tomato plants at 48 h (a) or 120 h (b) after inoculation with *Ralstonia* UW551 (orange), *Ralstonia* GMI1000 (blue), or water (gray). qRT-PCR was used to measure expression of class III acid, class III basic, and class II acid β-1,3-glucanase genes; expression levels are relative to those in similarly inoculated wilt-susceptible Money Maker plants at 48 or 120 hpi. (b and c) Wilt disease progress on wilt-susceptible MoneyMaker tomato of wild-type and _Δ_*acrA* and _Δ_*dinF* deletion mutants of *Ralstonia* UW551 (c) and *Ralstonia* GMI1000 (d). 17-day-old tomato seedlings were soil-soak inoculated at 5×10^7^ CFU/g of soil, and plants were rated on a 0 to 4 disease index scale. Symbols indicate mean disease index; results reflect 3 independent experiments, each with 15 plants per treatment. (e) Doubling time of *Ralstonia* UW551 wild type and _Δ_*acrA* and _Δ_*dinF* deletion mutants in *ex vivo* xylem sap from healthy or infected tomato plants as indicated in the legend. The wild-type strain grew similarly in all saps tested, but loss of either *dinF* or *acrA* slowed growth of UW551 in sap from resistant H7996 whether plants were infected by *Ralstonia* UW551 or *Ralstonia* GMI1000 (adjusted P<0.05 by ANOVA with Tukey’s multiple comparisons). Symbols indicate the average doubling times for 3 independent experiments, each containing 3 technical replicates; bars indicate standard error.

**Figure 6.**
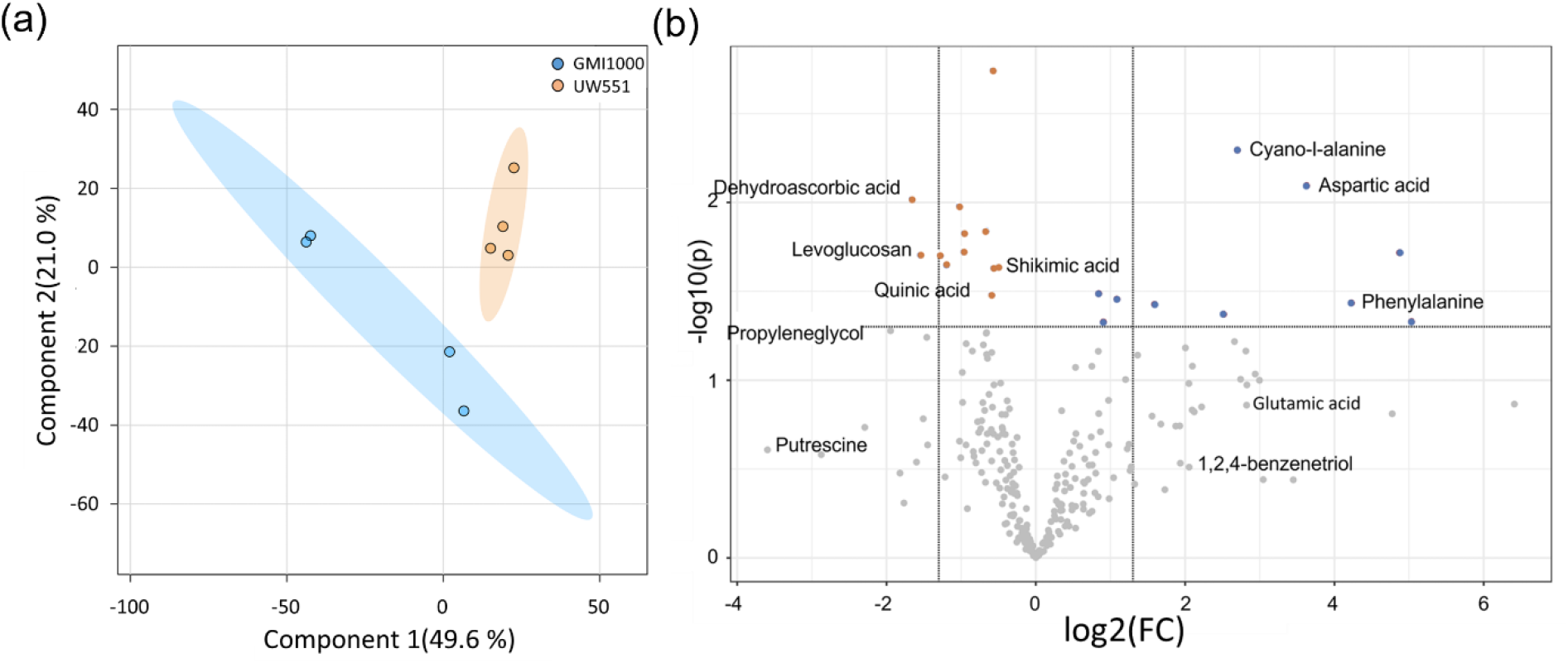
Metabolomic profiling of the xylem response of following Infection by *R. pseudosolanacearum* GMI1000 or *Ralstonia* UW551. (A) Two-dimensional principal component analysis (PCA) score plot illustrating the grouping of the variable conditions: Hawaii7996 ex vivo xylem sap harvested from plants infected with resistance-breaking UW551 (orange, H7U) and GMI1000 (blue, H7G). Ellipses on the score plot represent a 95% confidence interval. (B) Volcano plot combining results from Fold-Change (FC) Analysis and T-tests (p < .05) into a single graph to discern identified compounds based on biological interest, statistical significance, or both.

### 2.12 No single identified compound explains why xylem sap inhibits *Ralstonia* growth

We determined the minimum inhibitory concentration (MIC) of various differentially present compounds identified by the xylem sap metabolomic analysis, using the standard metric of qualitative bacterial growth in a twofold serial dilution of each tested compound. To control potential interactions of compounds with growth medium, each compound was tested following dilution in minimal medium supplemented with 0.2% glucose, rich CPG medium, and *ex vivo* xylem sap from healthy H7996 plants. Many tested compounds inhibited *Ralstonia* growth, but none differentially inhibited strain GMI1000 more than UW551 (Table 1). This result is consistent with our physical analysis suggesting that no single compound is likely to explain the inhibitory properties of H7996 xylem sap.

### 2.13 Pandemic lineage strains detoxify *ex vivo* xylem sap partially by reducing phenolic compounds

Plant chemical defenses typically include phenolic compounds, so the ability to degrade phenolics helps plant pathogens overcome host resistance. To determine if phenolic degradation affects *Ralstonia* strain growth in xylem sap, we measured total phenolic concentration in sap from healthy and infected plants. As expected, in response to *Ralstonia* infection total phenolic compounds increased in sap from both wilt-susceptible MM and resistant H7996 relative to sap from healthy plants. Strains UW551 and GMI1000 elicited similar phenolic levels in susceptible MM. However, in resistant H7996, total phenolics were much lower in response to UW551 than to GMI1000 (T-test, *P*< .05)(Fig 7A). To determine if these differences resulted solely from plant production or if plant-produced phenolics were degraded by the bacterium, we measured total phenolics in sap from *Ralstonia*-infected H7996 plants after it had been used as secondary growth medium for either GMI1000 (Fig 7B) or UW551 (Fig 7C). While both strains reduced phenolic levels somewhat, only sap that had been incubated with UW551 had significantly lower phenolic levels than untreated sap (Fig 7B-C) (adjusted *P*<0.05 by ANOVA with Tukey’s multiple comparisons). This indicated that greater ability to degrade defensive phenolic compounds in xylem sap could explain why *Ralstonia* strain UW551 overcomes H7996 wilt resistance.

**Figure 7.**
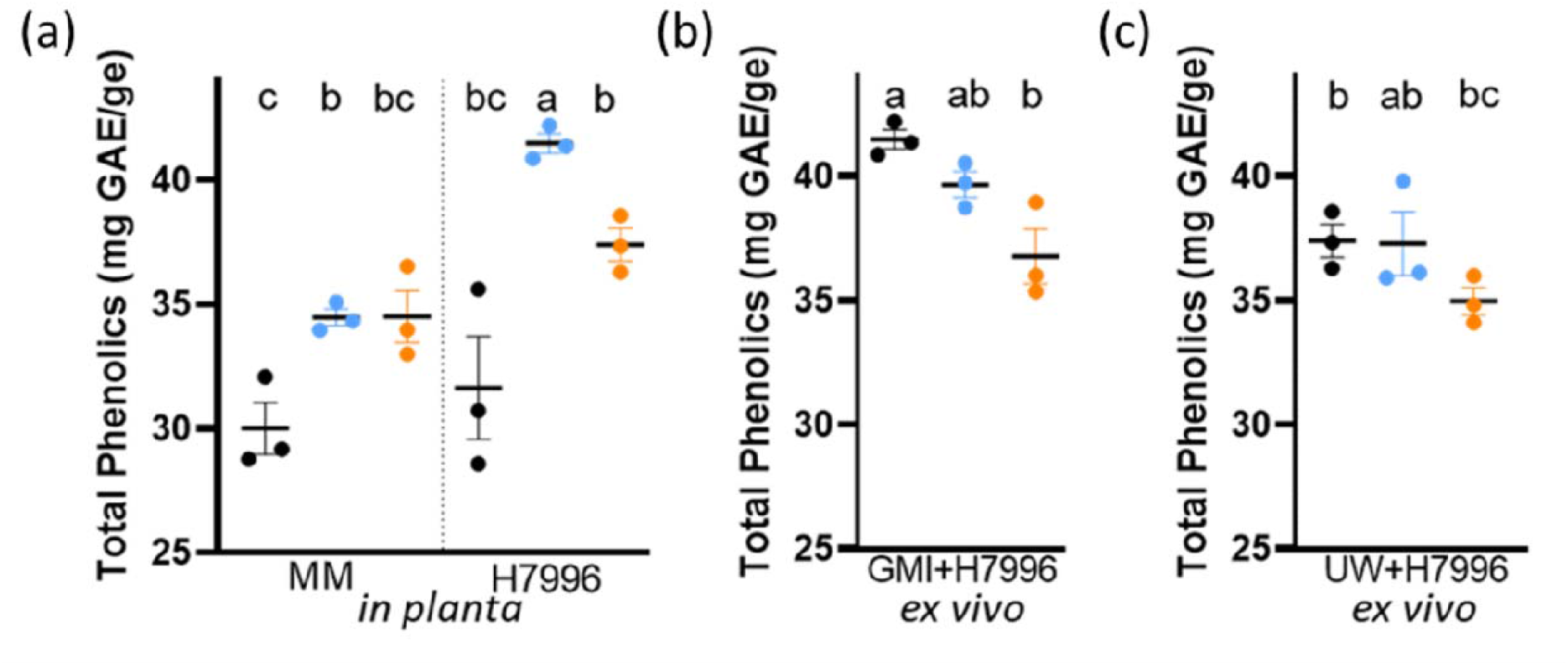
UW551 detoxifies ex vivo xylem sap partially by reducing phenolic compounds. (A) Total phenolic compounds in xylem sap harvested from wilt-susceptible Money Maker (MM) or resistant Hawaii 7996 (H7996) tomato plants infected with *R. pseudosolanacearum* GMI1000 (blue dots), *R. solanacearum* UW551 (orange dots) or water (black dots). Infection by either *Ralstonia* strain increased total phenolics in sap from both resistant and susceptible tomato plants relative to healthy plants (black dots). However, H7996 resistant plant sap contained more phenolics after GMI1000 infection (Blue dots) than after UW551 infection (orange dots) (B) Phenolic levels following 24 hours of growth of *Ralstonia* strain GMI1000 on sap harvested from H7996 plants infected *Ralstonia* GMI1000 (blue dots), *Ralstonia* UW551 (orange dots), or water (black dots). (C) Phenolic levels following 24 hours of growth of resistance-breaking *Ralstonia* strain UW551 on sap harvested from H7996 plants infected with *Ralstonia* GMI1000 (blue dots), *Ralstonia* UW551 (orange dots), or water (black dots) (adjusted P<0.05 by ANOVA with Tukey’s multiple comparisons). Each dot indicates the average of 3 technical replicates; each experiment contained 3 biological replicates.

## 3. Discussion

Most commercial tomato cultivars with bacterial wilt resistance trace all or part of their disease resistance to the Hawaii breeding lines, usually H7996 (Scott 2005). We found that *R. solanacerum* brown rot pandemic lineage strain UW551 overcomes H7996 resistance, causing wilting symptoms and killing around 40% of inoculated plants. This was a partial rather than a complete breakdown of H7996 resistance; in some cases UW551-inoculated plants recovered, which is a common characteristic of incomplete virulence (Jaworski et al., 1997; Grimault et al., 1995). Nevertheless, any threat to the durability of H7996-derived resistance warrants attention given the large global impact of bacterial wilt on tomato production (Mansfield et al., 2012).

It is commonly thought that host resistance to bacterial wilt acts primarily belowground, where most *Ralstonia* infections begin (Grimault 1994). Based on this idea, grafting agronomically desirable scions onto H7996 or other resistant rootstock is used to control bacterial wilt in fresh tomato production where the economic benefits of disease control outweigh the cost of grafting (McAvoy et al., 2012; Kunwar et al., 2015). In agreement with previous results, we found that grafting aboveground scions of H7996 to rootstocks of a susceptible tomato significantly reduced disease symptoms (Planas-Marquez 2020). This indicates that wilt disease resistance acts in the xylem of resistant tomato as well as in roots. Moreover, expression of JA-and SA-mediated host defense genes increased in susceptible rootstocks grafted to H7996 stems and in susceptible stems grafted to H7996 rootstocks. Using grafting to identify tissue-specific components of resistance showed at least part of this resistance comes from non-diffusible xylem structures and compounds, as has been described for other bacterial vascular diseases (Chatterjee et al., 2008). We conclude that in addition to root-specific responses, H7996 wilt resistance also functions in aboveground plant parts, including the xylem tissue colonized by *Ralstonia*.

Previous investigations of the tomato xylem environment have focused on susceptible cultivars or healthy plants (Chellemi et al., 1998; Zuluaga et al., 2013; Lowe-Power et al., 2018). A preliminary study of healthy tomato xylem sap chemistry identified differences in amino acid concentration between resistant and susceptible cultivars (Chellemi et al., 1998) and a comparative transcriptomic analysis of susceptible Ponderosa and resistant LS-89 tomato stems found that in response to *Ralstonia* infection, resistant plants increased expression of diverse defenses, including biosynthesis of phenolics and glucanases (Ishihara et al. 2012). A proteomic and gene silencing study found that GABA biosynthesis and the methionine cycle are required for full wilt resistance in H7996 plants, but not in susceptible controls; however, altering these elements of central metabolism likely has pleiotropic effects (Wang et al. 2019). By measuring biological outcomes rather than comparating biochemistry, we found that stems of H7996 co-inoculated with a mixture of GMI1000 and UW551 were more likely to be colonized by GMI1000 than when they were inoculated with GMI1000 alone. Although co-infection of a single plant by two or more *Ralstonia* strains has not been observed in the field, this experiment suggests that a resistance-breaking strain can make the host more susceptible to other microbes, including other *Ralstonia* strains. This is consistent with the finding that tomato plants infected by plant pathogenic Xanthomonads are better hosts for the food-borne human pathogen *Salmonella enterica* (Dixon et al., 2022).

Interestingly, bacterial growth on *ex vivo* xylem sap harvested from *Ralstonia*-infected plants recapitulates the whole plant response. We previously showed that *Ralstonia* infection makes *ex vivo* xylem sap from susceptible cultivars a better growth medium for the pathogen (Lowe-Power et al., 2018). Here, we found the opposite is true of resistant tomato cultivars; *Ralstonia* colonization created an unfavorable xylem environment. Specifically, *ex vivo* sap harvested from H7996 plants responding to inoculation by GMI1000, which does not wilt H7996, was poor growth medium for GMI1000. This result was generalizable to diverse other strains in the RSSC that also cannot overcome H7996 resistance. In parallel, phylotype III *Ralstonia* strain CMR15, which like UW55 can wilt H7996, also grew well in sap from infected H7996. However, another wilt-resistant tomato cultivar, CRA66, did not respond to *Ralstonia* infection by increasing inhibitors of bacterial growth in its xylem sap, suggesting that CRA66 resists bacterial wilt by a different mechanism than H7996. It would be interesting to determine if similar bacterial growth inhibitors are present in sap from wilt-resistant varieties of other *Ralstonia* hosts such as eggplant and tobacco.

Bacteria have many mechanisms for overcoming inhibitory environments (Brown et al., 2007; Deslandes and Rivas. 2012; Deslandes and Genin. 2014; Kim et al., 2016), suggesting several ways that UW551 might overcome H7995 resistance. Our experiments showed that neither carbon source availability, differential expression of β-1,3-glucanase genes, nor bacterial toxin efflux pumps could explain why xylem sap from H7996 inhibits growth of *Ralstonia* GMI1000 but not growth of resistance-breaking strain UW551. Stressors like *Ralstonia* infections provoke plants to relocate resources by mechanisms like induced lateral transport of sugars and inorganic solute restrictions (Aubry et al., 2019). However, total sugar content of resistant lines like H7996 does not differ from susceptible tomato (Georgelis and Scott, 2004). This aligns with our finding that reduced nutrient content does not explain the differential ability of GMI1000 and UW551 to grow in sap from previously infected H7996 plants.

A proposed mechanism of tomato bacterial wilt resistance is the differential expression of β-1,3-glucanases, enzymes involved in defensive callose deposition and degradation of plant cell wall polysaccharides (Levy et al., 2007; Ishihara et al., 2012). Although H7996 did indeed upregulate these enzyme genes much more than wilt-susceptible Money Maker in response to *Ralstonia* infection, none of the three tested classes of β-1,3-glucanases were differentially expressed in response to resistance-breaking strain UW551, suggesting that β-1,3-glucanases do not explain H7996 wilt resistance.

Bacteria can tolerate environmental toxins by expelling them with efflux pumps. We found that mutating *Ralstonia* efflux pumps AcrA and DinF reduced virulence of GMI1000 and UW551 on susceptible tomato as it did in *Ralstonia* strain K60 (Ravirala et al., 2007; Brown et al., 2007). Additionally, UW551ΔacrA and UW551Δdinf mutants trended towards slower doubling times in sap harvested from infected resistant plants relative to wildtype UW551 growth, although differences were not significant. The requirement for these pumps in sap suggest that in tomato xylem *Ralstonia* encounters sublethal levels of inhibitors like DNA-damaging compounds and hydrogen peroxide (Brown et al., 2007). It would be interesting, although technically challenging, to characterize the transcriptional response of an unsuccessful strain like GMI1000 to the toxic environment in H7996 xylem.

Eliminating these alternative explanations, together with the discovery that concentrating sap from GMI1000-infected H7996 plants made it even more inhibitory, suggested that induced chemical defenses inhibit unsuccessful *Ralstonia* strains. Although the precise antimicrobial effects of many plant-produced compounds are not yet defined, multiple mechanisms have been reported (Upadhyay et al., 2015). Our comparative metabolomics analyses found higher concentrations of many plant-derived antimicrobials in GMI1000*-*conditioned xylem sap compared to in sap from plants infected with UW551. These antimicrobial compounds inhibit bacteria by diverse mechanisms including leakage of cellular contents and loss of energy production (Tsuchiya et al., 2000). No known compound that was differentially present in our metabolomic analysis was more toxic to GMI1000 than to resistance-breaking strain UW551.

However, our minimal inhibitory concentration (MIC) assays were conducted using single compounds on bacteria grown in minimal medium; it is possible that *Ralstonia* strains respond differently in the plant environment to the combination of multiple compounds, especially potentially synergistic compounds in the same class. Strain UW551 did tolerate slightly higher concentrations of some plant-synthesized phenolic compound like caffeic acid. This might mean that plant compounds affect virulence in sub-lethal or sub-inhibitory concentrations as they do in another gram-negative pathogen, *Salmonella enteritidis* (Upadhyay et al., 2013). This aligns with the fact that QTL-mediated resistance is driven by multiple loci, often with overlapping functionality (Ye and Smith. 2008).

Resistance-breaking strain UW551 might grow better in stems of H7996 plants 1) because it does not trigger a strong defensive response; 2) because it can detoxify the resistant plant’s chemical defenses; or 3) both. Measuring pathogen growth in *ex vivo* xylem sap, in the absence of living plant tissue, allowed us to separate bacterial growth from plant defense. UW551 grew well in sap harvested from GMI1000-infected plants, but GMI1000 could not. However, this sap lost its inhibitory qualities after UW551 grew in it. This result shows that active detoxification of plant defenses explains at least part of UW551 success in H7996 plants.

Levels of phenolic compounds commonly increase during plant defense (Nicholson et al., 1992; Cai et al., 2008; Fortunato et al., 2012; Wang et al., 2020). Antimicrobial phenolics have been found in every tested *Ralstonia* host, but many don’t reach bactericidal concentrations (Nicholson et al., 1992). These include well-studied signaling molecules like salicylic acid (SA), which has a role in defense but can also accumulate to high concentrations with direct antimicrobial effects (Smith-Becker et al., 1998). Phenolics with high stability in aqueous systems have received the most attention as potential antimicrobial compounds. However, phenolic compounds range from low molecular weight phenols, such as the benzoic acids and the phenylpropanoids to high molecular weight polymerized phenols (Nagarajan et al., 2020). We previously observed reduced virulence in *Ralstonia* mutants unable to degrade salicylic acid or the trans-cinnamic acid derivatives ferulic acid and caffeic acid; these mutants were also more susceptible to inhibition by these compounds in culture (Lowe et al 2015, Lowe-Power et al 2016).

Xylem sap from H7996 plants infected with GMI1000 contained lower total phenolic compounds after UW551 grew in it than if GMI1000 was grown in it. This suggests that UW551 detoxifies xylem sap at least in part by degrading plant phenolic compounds. *Ralstonia* strains have several mechanisms for reducing phenolic compounds. Phylotype IIB strain IBSBF1503, which is related to UW551, increased expression of the Nag locus when IBSBF1503 was infecting melon rather than tomato (Ailloud et al., 2016). The Nag locus, which is also present in UW551, encodes the ability to degrade and detoxify SA. This could help *Ralstonia* detoxify the H7996 xylem environment. A transcriptomic analysis found that the oxygenase NagG is more strongly expressed in UW551 than in GMI1000 (absolute expression levels of 11.03 and 7.74 respectively) (Jacobs et al., 2012).

Previous metabolomic analyses demonstrate a consistent correlation between tomato bacterial wilt resistance and both preformed and induced phenolic compounds, notably trans-cinnamic acids (Shi et al. 2022; Zeiss et al 2022; Chen et al. 2021; Zeiss et al 2019; Zeiss et al 2018) add refs A recent physico-chemical study showed that infection of H7996 with Ralstonia GMI1000 triggered increased deposition of ligno-suberin and hydroxycinnamic acid derivatives on vessel walls, thereby restricting bacterial colonization (Kashyap 2022). Moreover, transgenically upregulating ligno-suberin production slowed disease progress on susceptible cv. Money Maker (Kashyap 2022). These elegant plant-side experiments provide complementary evidence that induced phenolic compounds are a key component of tomato resistance to bacterial wilt. It would be interesting to use similar physico-chemical techniques to determine if in addition to degrading soluble phenolics in xylem sap, resistance-breaking *Ralstonia* UW551 also degrades ligno-suberins and hydroxycinnamic acids deposited on vessel walls.

## Conclusions

In summary, metabolomic, chemical, and transcriptomic analyses, together with manipulative experiments using bacterial growth in xylem sap as a proxy for the plant environment, indicate that aboveground bacterial wilt disease resistance in breeding line H7996 is due at least in part to pathogen-induced antimicrobial phenolic compounds in xylem sap.

We identified two *Ralstonia* strains that break the widely-deployed H7996 wilt resistance: West African strain CMR15 and brown rot pandemic lineage strain UW551. Both these strains can detoxify inhibitory xylem sap. This result should inform plant breeders seeking to improve wilt resistance in tomato, especially varieties intended for the highland tropics where the *Ralstonia* brown rot pandemic lineage is present. Breeders should consider incorporating wilt resistance from other sources than H7996 and avoid resistance assays based only on root infection.

## 4. Experimental Procedures

### 4.1 Bacterial strains and culture

Bacterial strains used in this research are listed in Supplementary Table S3. *Ralstonia* strains were routinely cultured in either minimal BMM medium or rich CPG medium (Khokhani et al. 2018). Bacterial growth was measured spectrophotometrically as optical density at 600 nm (OD_600_), using volumes of 1 ml, 200 μl, or 50 μl.

### 4.2 Tomato plant growth and grafting

Tomato cultivars used in this research are listed in Supplementary Table S3. Plants were sown, transplanted, and grown as described (Khokhani et al. 2018). Fully expanded first true leaves were present 24-48 h sooner on H7996 wilt resistant tomato plants than on wilt-susceptible Bonny Best or Moneymaker tomato plants, so H7996 seedlings were transplanted sooner to ensure all plants were at a comparable growth stage. Tomato plants were grafted when rootstock plants had five true leaves (∼28 days after sowing), and the scion plants had three true leaves (∼21 days after sowing) (Kunwar et al 2015).

### 4.3 Construction of *Ralstonia* mutants

Cloning, restriction digestion, sequencing, and PCR used standard methods (Castaneda et al. 2005; Heckman and Pease 2007; Monteiro et al. 2012). Gibson assembly was used to create constructs with complete in-frame deletions of the gene(s) of interest, using the primers listed in Supplementary Table S3, and cloned into the suicide vector pstBlue with gentamcin or kanamycin cassette insertion (Schweizer, 1993). Deletion constructs were introduced into the chromosome of wild-type *Ralstonia* GMI1000 or UW551 via double recombination to create mutant strains GMI1000ΔacrA, GMI1000ΔdinF, UW551ΔacrA, and UW551ΔdinF. Following electroporation double recombinants were selected on 15mg/L gentamicin and 25 mg/L kanamycin. Correct allelic replacement for each mutant strain was confirmed by PCR and DNA sequencing. DNA sequencing was performed at the University of Wisconsin-Madison Biotechnology Center and oligonucleotide primers were synthesized by Integrated DNA Technologies (Coralville, Iowa).

### 4.4 Tomato xylem sap collection

The tomato xylem sap harvest process attempted to minimize variability in colonization, flow rate, and plant physiology. Sap was harvested from well-watered tomato plants 5 days after inoculation with *Ralstonia* or water (healthy control) and within 1-4 hours after beginning of the growth chamber light period (Khokhani et al. 2018; Lowe-Power et al. 2018), with modifications. Briefly, after plants were detopped 2 cm above the cotyledons, root pressure forced xylem sap to accumulate on the stump. After 5 min, the stump was washed with water to remove debris and cytoplasm from broken cells. Over the next 60 min, sap accumulating on the stump was frequently collected with a pipette and transferred into pre-chilled 96 well 22-micron filter plates on ice or a 4°C block. Plates were centrifuged at 4,000 rpm at 4°C for 4 min to filter-sterilize the *ex vivo* xylem sap, which was flash-frozen and stored at -80°C until analysis. For each plant from which xylem sap was harvested, the population size of *Ralstonia* in the stem was measured by serial dilution plating as described below under colonization assays. Only sap from plants containing between 5 × 10^4^ and 5 × 10^8^ CFU per gram of stem was pooled and retained for future experiments. Individual sap samples were pooled across sampling times and biological replicates as described (Lowe-Power et al 2018).

### 4.5 Bacterial growth in xylem sap

We measured *Ralstonia* growth in *ex vivo* xylem sap as a culture medium (Hamilton et. al. 2021). Briefly, *Ralstonia* cells were inoculated into 50 μl of pooled, filter-sterilized xylem sap in 96-well plates. Plates were incubated for 24 h in a BioTek Plate reader and growth was measured spectrophotometrically as O.D._600_ and quantified as the area under the growth curve. The Mann-Whitney test was used to analyze the growth data relative to growth of the same *Ralstonia* strain in xylem sap from healthy plants.

### 4.6 GC-MS metabolomics of xylem sap

*Ex vivo* xylem sap from tomato plants was subjected to untargeted metabolomic analysis using GC-MS. Sap from healthy H7996 tomato plants was compared to sap from plants that had been soil-soak inoculated with *Ralstonia* strain GMI1000 or UW551 and verified to contain >5×10^4^ CFU g^-1^ stem. Sap samples harvested as described above were thawed and pooled across sampling time and 4 biological replicates. Five plant samples were pooled, flash frozen, shipped on dry ice to West Coast Metabolomics (U. California-Davis), and characterized using their untargeted metabolomic pipeline. West Coast Metabolomics did preliminary data analysis including peak noise filtering, metabolite quantification by peak height, and metabolite annotation. We normalized metabolite peak heights by sample final volume for each of the four pooled sample because exudation rate alters sap metabolite concentration (Goodger et al., 2005). In some cases, significantly different metabolites had FDR <0.1. Statistical analyses were conducted in MetaboAnalyst 5.0 using the previously described pipeline (Lowe-Power et al 2018).

### 4.7 Measuring phenolic compounds in xylem sap

The total phenolic content of xylem sap samples was determined using the Folin-Ciocalteu assay (Singleton et al 1999). This electron transfer-based method measures reducing capacity, which is expressed as phenolic content. External calibration was done using known concentrations of gallic acid. Reaction absorbance was measured spectrophotometrically at 750 nm. Total phenolic content of each sample was calculated as mM gallic acid equivalent (mM GAE) by extrapolation from a gallic acid standardization curve.

### 4.8 Determination of Minimum Inhibitory Concentrations

The MIC of a chemical was defined as the lowest concentration that prevents detectable cell growth as determined by spectrophotometric comparison to an uninoculated control. MIC assays were carried out in 96-flat-bottomed well microtiter plates using as growth medium either BMM supplemented with 0.2% glucose or *ex vivo* xylem sap harvested from healthy tomato plants (described above and hereafter described as “healthy sap”). Growth media were supplemented with different compounds at various concentrations. Each well received 100 μl (for BMM) or 25μl (for xylem sap) of a twofold dilution series of an inhibitor solution and 100 μl or 25μl of bacterial suspension. All inhibitor solvents were included as controls for effects on bacterial growth.

### 4.9 Virulence assays

Tomato plants were inoculated using a naturalistic soil-soak (Khokhani et al. 2018). Briefly, 50 ml of a water suspension of bacteria was poured onto the soil of unwounded 17-day-old tomato plants growing in 4-inch pots in a 28°C growth chamber to achieve a final pathogen density of 5 × 10^7^ CFU /gm soil. Plants were rated daily for symptoms using a 0 to 4 Disease Index scale. All experiments were repeated at least three times with 15 plants/treatment/experiment. Virulence data were analyzed using repeated measures analysis of variance (RM-ANOVA), which is used for independent variables that are related because they are sampled repeatedly over time.

### 4.10 Measuring bacterial colonization of plants and competitive fitness

Plants were directly inoculated with *Ralstonia* via a cut petiole and bacterial colonization of stems was quantified (Khokhani et al. 2018). Briefly, 10 μl suspension of 2,000 bacterial cells was applied to a freshly-cut leaf petiole. To assess strain competitive fitness, plants were inoculated with a 1:1 ration of antibiotic resistance–marked wild type and mutant cells. All experiments were repeated at least three times with 15 plants/treatment/experiment. Data were analyzed using a Mann-Whitney *U* test or Wilcoxon sign-rank test.

### 4.11 Quantitative real-time PCR (qRT-PCR) analysis

Samples were taken from mid stem and roots of tomato plants and total RNA was extracted using a hot-phenol chloroform method (Jacobs et. al 2012). For each sample, 2 μg of total RNA was reverse transcribed into cDNA using SuperScript III cDNA Synthesis kit (ThermoFisher) according to the manufacturer’s instructions. Subsequent qRT-PCR reactions were performed using the primers in Supplementary Table S3 (Block et. al 2005). Relative gene expression compared to untreated control plants was determined using the ΔΔCT method (Livak & Schmittgen, 2001).

## Acknowledgements

This research was supported by the University of Wisconsin-Madison College of Agricultural and Life Sciences. CDH was supported by a USDA-NIFA Predoctoral Fellowship & a UW-Madison SciMed Graduate Research Scholarship. The authors thank Dr. Sanju Kunwar (U. Florida), Dr. Tiffany Lowe-Power (U. California-Davis), Dr. Anjali Iyer-Pascuzzi (Purdue U.), Dr. Jessica Cooperstone (Ohio State U.), Dr. Alicia N. Truchon and Mariama D. Carter (U. Wisconsin-Madison) for technical advice and helpful discussions.

## Data availability statement

Links to metabomics datasets are given in Supplemental Table S1. Other data that support the findings of this study are available from the corresponding author upon reasonable request.

## Table Legend

Table 1. Minimal inhibitory concentration (MIC) of selected compounds (in μg/ml) for *Ralstonia* strains GMI1000 and resistance-breaking UW551

## Supporting Information Legends

**Supplemental Figure 1. Tomato wilt resistance or susceptibility shapes *Ralstonia* growth in *ex vivo* xylem sap**. (A) Growth of *Ralstonia* strain (indicated on X-axis) on filter-sterilized xylem sap harvested from tomato cultivars (indicated on the left Y-axis) that were infected by various *Ralstonia* strains as indicated on the right Y-axis. (B) Growth of diverse *Ralstonia* strains on filter-sterilized xylem sap harvested from wilt-resistant H7996 and wilt-susceptible Money Maker tomato plants (indicated on right Y-axis) that were either healthy or infected with various *Ralstonia* strains as indicated on the left Y-axis. For all *ex vivo* sap experiments, bacterial growth was measured in a plate reader over 24 h spectrophotometrically as A_600nm_. Bacterial growth, quantified as area under a 24-hour growth curve, is indicated by shading from white (low area or little to no growth) to black (maximum growth). Data are from 3-6 independent experiments, each with 3-6 technical replicates.

**Supplemental Figure 2. Overview of xylem sap metabolomic characterization**. Filter-sterilized *ex vivo* xylem sap harvested from Hawaii7996 and Money Maker tomato plants following infection by *R. pseudosolanacearum* GMI1000 or *Ralstonia* UW551; sap from healthy plants served as controls. (A) A 2D principal component analysis (PCA) score plot grouping the variable conditions: healthy Hawaii7996 (green, H7H), healthy Money Maker (purple, MMH), Infected Hawaii7996 (orange, H7+) and infected Money Maker (blue, MM+). (B) PCA score plot of compounds identified in sap from H7996 plants that were healthy (grey), or infected by UW551 (orange) or GMI1000 (blue). The ellipse on the score plot represents a 95% confidence interval. (C) Top 25 VIP (variable importance in projection) scored compounds identified in *ex vivo* xylem sap from H7996 plants that were healthy (H7H) or infected by UW551 (H7U) or GMI1000 (H7G). In a Partial Least Squares Discriminant Analysis (PLS-DA) the VIP score is used to distinguish samples. The colored boxes on the right indicate the relative concentrations of the corresponding metabolite in each group under study. (D) A heatmap that provides intuitive visualization of UW551 and GMI1000 conditioned Hawaii7996 ex vivo xylem sap. Each colored cell on the map corresponds to a concentration value, with metabolites in rows and samples in columns. The heatmap identified top 25 metabolites that are unusually high/low relative to healthy Hawaii7996 ex vivo xylem sap.

**Supplemental Table 1**. Links to metabolomic data sets

**Supplemental Table 2**. Methods for characterizing bacterial growth inhibitor

**Supplemental Table 3**. Plant pathogenic *Ralstonia* strains, plant cultivars, and primers

## Notes

### Competing Interest Statement

The authors have declared no competing interest.

