## Supplemental Materials for "*Ralstonia solanacearum* pandemic lineage strain UW551 overcomes inhibitory xylem chemistry to break tomato bacterial wilt resistance"

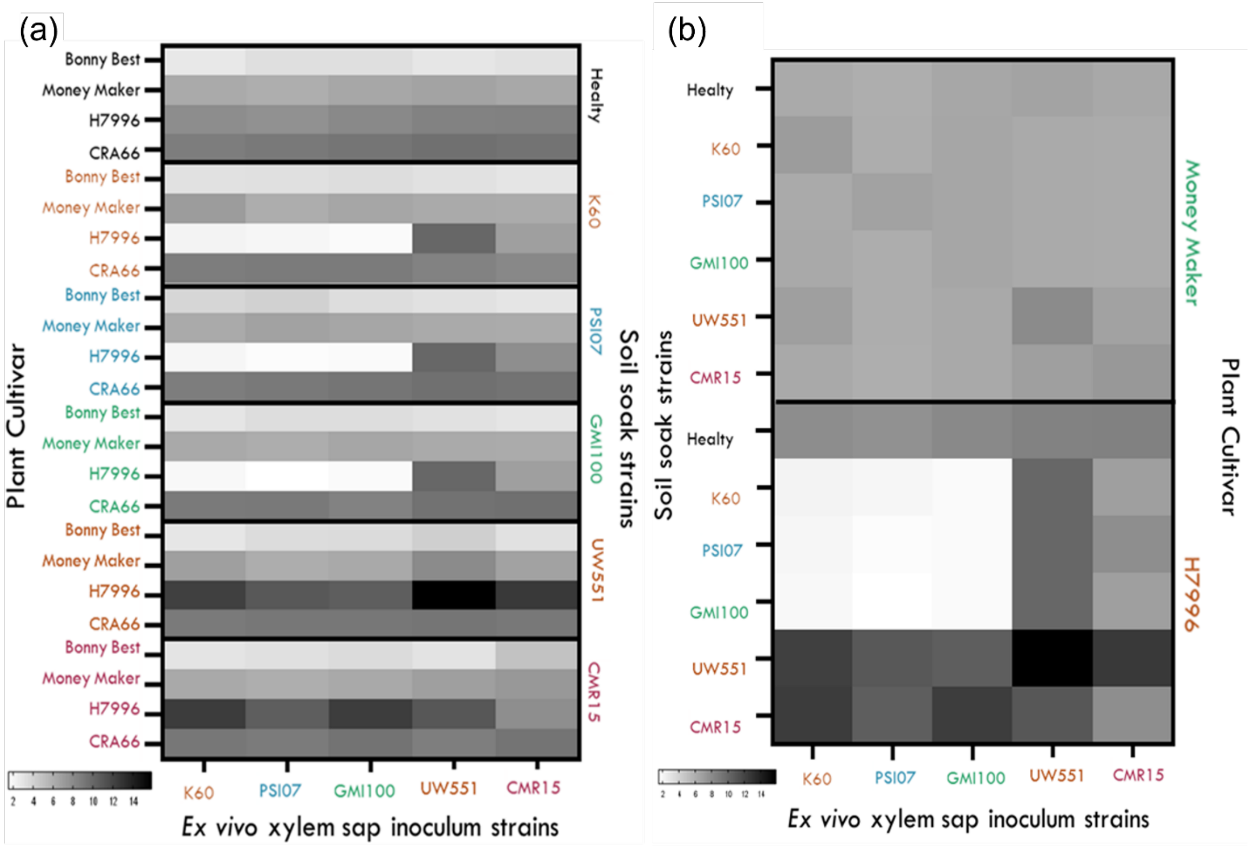


**Supplemental Figure 1. Tomato wilt resistance or susceptibility shapes *Ralstonia* growth in *ex vivo* xylem sap.** (a) Growth of *Ralstonia* strain (indicated on X-axis) on filter-sterilized xylem sap harvested from tomato cultivars (indicated on the left Y-axis) that were infected by various *Ralstonia* strains as indicated on the right Y-axis. (b) Growth of diverse *Ralstonia* strains on filter-sterilized xylem sap harvested from wilt-resistant H7996 and wilt-susceptible Money Maker tomato plants (indicated on right Y-axis) that were either healthy or infected with various *Ralstonia* strains as indicated on the left Y-axis. For all *ex vivo* sap experiments, bacterial growth was measured in a plate reader over 24 h spectrophotometrically as $A_{600nm}$. Bacterial growth, quantified as area under a 24-hour growth curve, is indicated by shading from white (low area or little to no growth) to black (maximum growth). Data are from 3-6 independent experiments, each with 3-6 technical replicates.

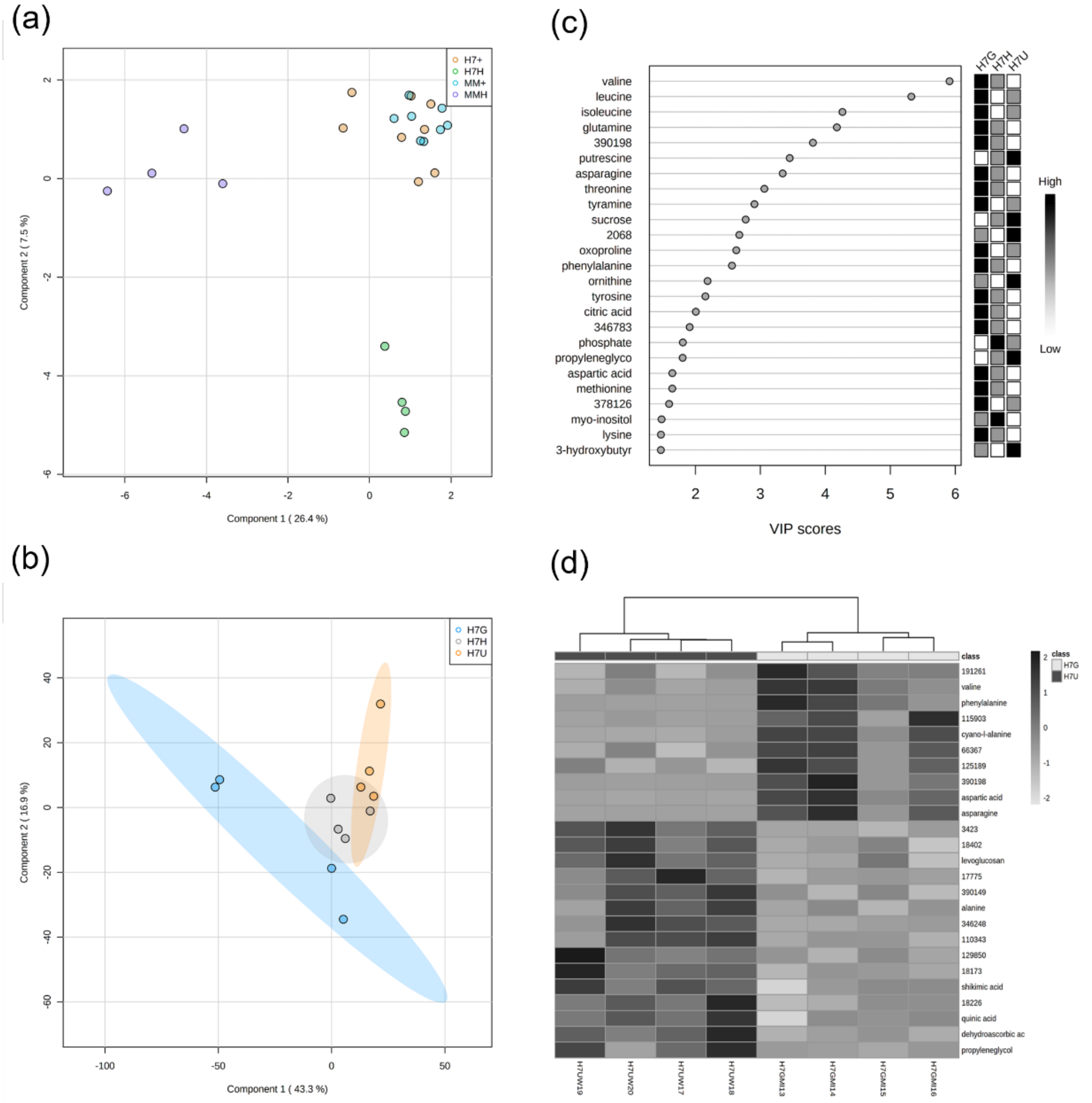


**Supplemental Figure 2:** Overview of xylem sap metabolomic characterization. Filter-sterilized *ex vivo* xylem sap harvested from Hawaii7996 and Money Maker tomato plants following infection by *R. pseudosolanacearum* GMI1000 or *Ralstonia* UW551; sap from healthy plants served as controls. (a) A 2D principal component analysis (PCA) score plot grouping the variable conditions: healthy Hawaii7996 (green, H7H), healthy Money Maker (purple, MMH), Infected Hawaii7996 (orange, H7+) and infected Money Maker (blue, MM+). (b) PCA score plot of compounds identified in sap from H7996 plants that were healthy (grey), or infected by UW551 (orange) or GMI1000 (blue). The ellipse on the score plot represents a 95% confidence interval. (c) Top 25 VIP (variable importance in projection) scored compounds identified in *ex vivo* xylem sap from

H7996 plants that were healthy (H7H) or infected by UW551 (H7U) or GMI1000 (H7G). In a Partial Least Squares Discriminant Analysis (PLS-DA) the VIP score is used to distinguish samples. The colored boxes on the right indicate the relative concentrations of the corresponding metabolite in each group under study. (d) A heatmap that provides intuitive visualization of UW551- and GMI1000-conditioned Hawaii7996 *ex vivo* xylem sap. Each colored cell on the map corresponds to a concentration value, with metabolites in rows and samples in columns. The heatmap identified top 25 metabolites that are unusually high/low relative to healthy Hawaii7996 ex vivo xylem sap.

**Supplemental Table 1: Links to metabolomic datasets**

| Name | Data Sheet Link | Contents |
|---|---|---|
| MetabolomicsAll | https://docs.google.com/spreadsheets/d/1ZqSys9fkt6h3jfZGz | Experimental control with pooled samples |
| MetabolomicsMM | https://docs.google.com/spreadsheets/d/1ZqSys9fkt6h3jfZGz | Cultivar controls |
| MetabolomicsH7 | https://docs.google.com/spreadsheets/d/1ZqSys9fkt6h3jfZGz | PCA plots & Fold-change analysis |
| MetabolomicsGMI1000 | https://docs.google.com/spreadsheets/d/1ZqSys9fkt6h3jfZGz | Fold-change analysis |
| MetabolomicsUW551 | https://docs.google.com/spreadsheets/d/1ZqSys9fkt6h3jfZGz | Fold-change analysis |

**Supplemental Table 2.** Methods for characterizing bacterial growth inhibitor

| Manipulation Method | Treatment Selection | Compound Characteristics[a] |
|---|---|---|
| Dehydration (Speed Vac) | Excludes volatiles | Stable aromatics |
| Heat (65°C for 15 mins) | Denaturation | Heat intolerant |
| Proteinase treatment | Degrades proteins | Protein-containing |
| Freezing (-80°/20°C) | Cold storage survival | Cold tolerant |
| Proteus X-Spinner | Size exclusion | Larger than 20kDa |
| Phase Lock Tubes | Separates organic and water-soluble compounds | Organic |

[a]General trait classification based on manipulation treatment (see Methods for details)

**Supplemental Table 3:** Plant pathogenic *Ralstonia* strains, plant cultivars, and primers

| Strains | Description[a] | Primers[bc] | Reference[d] |
|---|---|---|---|
| GMI1000[e]<br><br>(RUN054, JS753, CFBP6424. UW643) | *Ralstonia pseudosolanacearum* phylotype I seq 18 | N/A | Digat, 1978 |
| GM1000ΔacrA | *acrA* deletion mutant<br><br>RND family efflux pump | Up<br><br>F - tgcagacgcgttacgtatcggaagttggagcgggtcttg<br><br>R - actgagcgtcctgaaagccggtgacaag<br><br>Kan[r]<br><br>F - cggctttcaggacgctcagtggaacgaaaac<br><br>R - ccatccgcaatcatgaacaataaaactgtctgcttac<br><br>Down<br><br>F - ttgttcatgattgcggatggcgttgtag<br><br>R - gaatacagcggccgcgagctgccgcagtaaacaaggag | This study |

| | | | |
|---|---|---|---|
| GM1000ΔdinF | *dinF* deletion mutant MATE family efflux pump | Up<br><br>F - ttgttcatgatttatcgatcgcccggtc<br><br>R - gaatacagcggccgcgagcttcgtcgagcttggccttg<br><br>$Gm^r$<br><br>cctccacagatcatgtgcagctccatcag<br><br>agtcgtgatctgagctcagccaatcgac<br><br>Down<br><br>F - tgcagacgcgttacgtatcgcgtctgaccatgatgccc<br><br>R - actgagcgtcgcaaacgaatgcaccacc | This study |
| UW551[e] | *Ralstonia solanacearum*; Phylotype IIB seq 1<br><br>Brown rot pandemic lineage | N/A | Williamson, 2002 |

| | | | |
|---|---|---|---|
| UW551ΔacrA | *acrA* deletion mutant<br>RND family efflux pump<br><br>Gent$^R$ | Up<br>F - ttgttcatgatttatcgatcgcccggtc<br>R - gaatacagcggccgcgagcttcgtcgagcttggccttg<br>Kan$^R$<br>F - attcgtttgcgacgctcagtggaacgaaaac<br>R - gatcgataaatcatgaacaataaaactgtctgcttac<br>Down<br>F - tgcagacgcgttacgtatcgcgtctgaccatgatgccc<br>R - actgagcgtcgcaaacgaatgcaccacc | This study |
| UW551ΔdinF | *dinF* deletion mutant MATE family efflux pump<br>Gent$^R$ | Up<br>F - gctgagctcagatcacgactgcaccgatg<br>R - gaatacagcggccgcgagctcaacctgcaccgatttgg<br>GM$^r$<br>F - cctccacagatcatgtgcagctccatcag<br>R - agtcgtgatctgagctcagccaatcgac<br>Down<br>F - tgcagacgcgttacgtatcgccaacaatacggcttgtg<br>R - ctgcacatgatctgtggaggtgttgcttc | This study |

| | | | |
|---|---|---|---|
| PSI07[e]<br><br>(UW658, RUN083) | *Ralstonia syzgii*; phylotype IV seq | N/A | (Prior, 2005) |
| K60[e]<br><br>(UW25, CFBP2047) | *Ralstonia solanacearum*; Phylotype IIA seq 7 | N/A | (Collection by Kelman; 1949) |
| CMR15[e]<br><br>(RUN133, CFBP6941, UW663) | *Ralstonia pseudosolanacearum*; phylotype III seq 36 | N/A | (Prior, 2005) |
| **Tomato Genotypes** | **Bacterial Wilt Response** | **Primers** | **Reference** |
| Hawaii7996 | Resistant | N/A | Gilbert et al., 1973 |
| Money Maker | Susceptible | N/A | |
| CRA66 | Resistant | N/A | Prior et al., 1994 |
| Bonny Best | Susceptible | N/A | |
| **qPCR Primers** | **Description** | **Primers** | **Reference** |

| | | | |
|---|---|---|---|
| Pr-1a | Marker of salicylic acid (SA) defense signaling pathway | F - GAGGGCAGCCGTGCAA<br><br>R - CACATTTTTCCACCAACACATTG | Milling et al., 2011 |
| PR-1b | Marker of ethylene (ET) defense signaling pathway | F - TTGGTGACTGCGGGATGA<br><br>R - GGCGGCGGCTAGGTT T | Milling et al., 2011 |
| Pin2 | Marker of jasmonic acid (JA) defense signaling pathway | F - TGATGCCAAGGCTTGTACTAGAGA<br><br>R - AGCGGACTTCCTTCTGAACGT | Milling et al., 2011 |
| TomQ'a | β-1,3-glucanase<br><br>class III acidic | F - AAGCAAGAAGAGAGCATTAAAAGG<br><br>R -GTAATATGTTGGTTTCTTTATTAGCATATG | Ishihara et al., 2012 |
| PRQb | β-1,3-glucanase<br><br>class III basic | F - ACGCGTTGTTTACATCCCCTGGA<br><br>R - AGTTGTTGTTGTAAGTCCTCGCGT | Ishihara et al., 2012 |
| A-glu | β-1,3-glucanase<br><br>class II acidic | F - AACAGGAGCGCAGCCTATCGG<br><br>R - CCTTGGCGTTTGGAAGGATTGGC | Ishihara et al., 2012 |
| Actin | consistently expressed gene for qRT-PCR normalization | F - TCAGCAACTGGGATGATATG<br><br>R - TTAGGGTTGAGAGGTGCTTC | Milling et al., 2011 |
| *DnaJ*-like | consistently expressed gene for qRT-PCR normalization | F - ATGAAGCGCCAGATACCATC<br><br>R - TCAAGGCTCAATGTGTGCTC | Milling et al., 2011 |

| | | | |
|---|---|---|---|
| UBI3 | consistently expressed gene for qRT-PCR normalization | F- AGGTTGATGACACTGGAAAGGTT<br><br>R - AATCGCCTCCAGCCTTGTTGTA | Ishihara et al., 2012 |

[a] Marker genes for activation of defense signaling pathways: JA- jasmonic acid, ET – ethylene, SA – salicylic acid. Actin, *DnaJ*-like and UBI3 expression levels were used for normalization.

[b] Primers were designed using the primer3 program from Biology Workbench. Primer specificity for the corresponding gene was confirmed by BLAST search ( www.ncbi.nlm.nih.org).

[c] Uppercase primers were designed for qPCR while lowercased primers were designed for creating Gibson assembly cloning fragment to move from parent strain and pRCkan$^r$ or pRCGm$^r$ into pstBlue transformation ready plasmid.

[d] Reference publication and parenthesis indicate collector and year of collection of isolates used in this study.

[e] Name of strain as used in this study; names in parenthesis are alternative strain names.
